# Neural circuit analysis using a novel intersectional split intein-mediated split-Cre recombinase system

**DOI:** 10.1101/2020.04.25.061341

**Authors:** Audrey Tze Ting Khoo, Paul Jong Kim, Ho Min Kim, H. Shawn Je

## Abstract

The defining features of a neuron are its functional and anatomical connections with thousands of other neurons in the brain. Together, these neurons form functional networks that direct animal behaviour. Current approaches that allow the interrogation of specific populations of neurons and neural circuits rely heavily on targeting their gene expression profiles or connectivity. However, these approaches are often unable to delineate specific neuronal populations. Here, we developed a novel intersectional split intein-mediated split-Cre recombinase system that can selectively label specific types of neurons based on their gene expression profiles and structural connectivity. We developed this system by splitting Cre recombinase into two fragments with evolved split inteins and subsequently expressed one fragment under the influence of a cell type-specific promoter in a transgenic animal, and delivered the other fragment via retrograde viral gene transfer. This approach results in the reconstitution of Cre recombinase in only specific population of neurons projecting from a specific brain region or in those of a specific neuronal type. Taken together, our split intein-based split-Cre system will be useful for sophisticated characterization of mammalian brain circuits.

## Introduction

The human brain is arguably the most complex and sophisticated organ made up of intricate networks of neurons connected to each other across different brain regions. These neuronal networks process external stimuli, produce sensory and emotional experiences to generate calculated responses to the environmental stimuli based on intrinsic mechanisms or learned experiences (McIntosh, 2000; Schultz, 2000; Sporns, 2013). A fundamental goal in neuroscience therefore is to understand the functions of individual neurons and to delineate how different groups of neurons function together to orchestrate animal behavior (Yuste, 2015). To achieve this goal, researchers have utilized molecular and genetic tools in model organisms using binary expression systems, such as the Cre-LoxP system in mice and the GAL4-UAS system in *Drosophila* (del Valle Rodríguez et al., 2012; Tsien, 2016). Coupled with recent technological advances, including optogenetics to manipulate neuronal activity, voltage- or calcium-sensing proteins to visualize activity, and genetically engineered animals, these binary expression systems have tremendously expanded our knowledge of certain neuronal populations (Tsien, 2016; Wyart et al., 2009).

Although the Cre-LoxP system has been vital in helping us understand neuronal function, a single gene expression profile is often insufficient to target specific neuronal populations of interest (Hirrlinger et al., 2009; Jullien et al., 2003). For example, recent results from single-cell RNA sequencing analyses indicate high levels of transcriptomic heterogeneity among neurons of the same subtype, even within the same brain subregions (Lake, 2016). As such, intersectional bipartite systems were developed to target more genetically defined and homogenous populations of neurons. Hirrlinger et al. (2009) utilized a Cre-based complementation system by splitting Cre recombinase into two complementation-competent Cre protein fragments, allowing expression of the two split-Cre fragments to be driven by two different promoters. This strategy limited the expression of Cre recombinase to only cells that expressed both fragments of Cre (Hirrlinger et al., 2009). Similarly, Jullien et al. (2007) split Cre recombinase into two complementary-competent fragments that could be reconstituted via a ligand to achieve better temporal control over the expression of Cre recombinase. Despite their utilities, these split-Cre systems have not been widely implemented because Cre recombination depended on the proximity of the two split-Cre fragments, thus making it challenging to ensure reliable and stable association and specific nucleotide cleavage within the targeted DNA. Here, we present an improved method to reconstitute Cre recombinase using split inteins, which are self-catalytic protein elements that facilitate protein trans-splicing reactions (Lockless and Muir, 2009; Perler, 2002; Shah, 2011). This approach is therefore advantageous over other split-Cre methods that require exogenous ligands for recombination (Zheng et al., 2012). By using this split intein-mediated split-Cre recombinase system, we aimed to label long-range GABAergic projection neurons that could not be genetically targeted with the current research tools (Jinno et al., 2007; Lee et al., 2014; Margolis et al., 2006; Melzer et al., 2017). By simply expressing one split-Cre fragment in the neurons of the GABAergic lineage and delivering the other fragment via retrograde viral gene transfer, we could constitute Cre activity in only long-range GABAergic neurons that projected their axons from the central amygdala (CeA) to the dorsal striatum (DS).

## Results

We split Cre recombinase into two fragments, NCre (aa1-59) and CCre (aa60-343) (Jullien et al., 2003), and attached an N- or C-intein of the evolved split intein Npu37, which can express and trans-splice efficiently in mammalian cells (Lockless and Muir, 2009), to the C-terminal sequence of NCre and the N-terminal sequence of CCre respectively. This resulted in two fusion genes: NCre-IntN and IntC-CCre (Fig. 1A). The transcription and translation of the NCre-IntN and IntC-CCre fusion genes lead to the binding and autocatalytic trans-splicing of the split inteins. This reaction then ligates NCre and CCre together via peptide bonds to form functional Cre recombinase with additional peptide sequences (KGCFNKEDGS from IntC and GFL from IntN) (Shah and Muir, 2014). Based on our 3D modeling, the additionally incorporated peptide sequences do not occlude the active DNA-binding site of the reconstituted Cre recombinase (Fig. 1B). To test whether split intein-mediated trans-splicing could occur in mammalian cells under physiological conditions, we expressed epitope-tagged NCre-IntN and IntC-CCre constructs in HEK293T cells, and the resulting lysates were subjected to western blot analysis (Fig. 1C and D). When either NCre-IntN or IntC-CCre was transfected into HEK293T cells alone, only a single 49 kD band or a 37 kD band was observed by western blot using specific antibodies against either HA or FLAG respectively (Fig. 1D). However, when both NCre-IntN and IntC-CCre were transfected, we observed a band of higher molecular weight (67kD) detectable by both HA and FLAG specific antibodies, indicating successful split intein-mediated trans-splicing reaction under physiological conditions (Fig. 1D).

**Figure 1.**
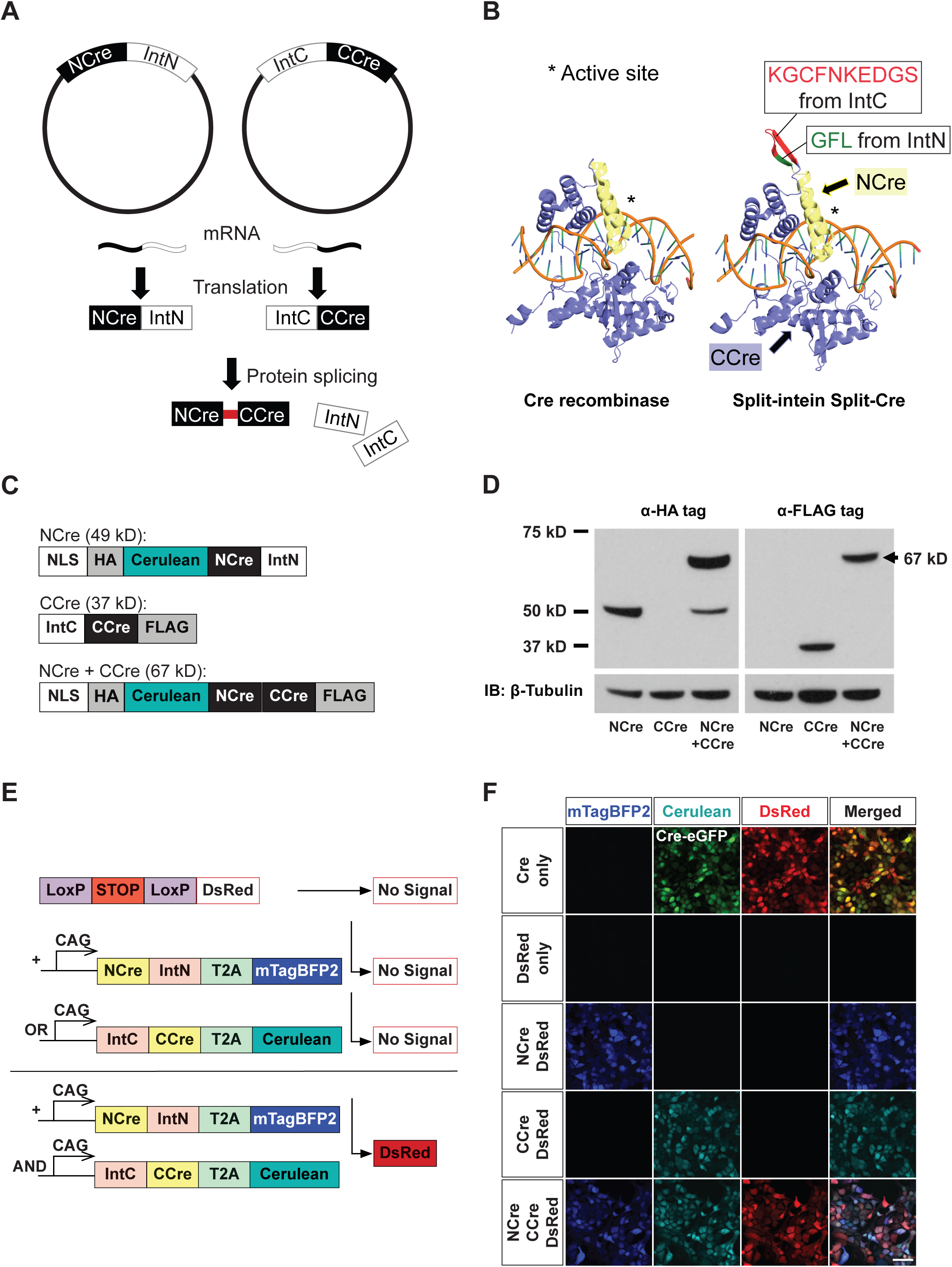
Design of the split intein-mediated split-Cre recombinase system. 1(A) Schematic depicting the autocatalytic split intein-mediated trans-splicing reaction to reconstitute Cre recombinase. Split inteins associate to fuse NCre and CCre with a peptide bond. 1(B) Structural comparison of original Cre recombinase-LoxP DNA complex (left, PDB ID: 1NZB (Ennifar et al., 2003) with a model for split intein-mediated split-Cre recombinase-LoxP DNA complex (right). The structure of split inteins from IntN (GFL) and IntC (KGCFNKEDGS), NCre and CCre are colored as green, red, yellow and blue respectively. Conserved active site are marked with an asterisk. 1(C) Schematic demonstrating the predicted size of the protein when NCre-IntN tagged with HA (49 kD) or IntC-CCre tagged with FLAG (37 kD) are transfected alone. A 67 kD reconstituted product is expected when both NCre-IntN and IntC-CCre are cotransfected. 1(D) Western blot analysis showing that when NCre-IntN is transfected alone, a 37 kD protein band was observed when western blotting was performed using a specific antibody against HA. When IntC-CCre was transfected alone, a 49 kD protein band was observed when western blotting was performed using a specific antibody against FLAG. When NCre-IntN and IntC-CCre were cotransfected, a 67 kD protein band with a higher molecular weight was observed when western blotting was performed using a specific antibody against HA or FLAG. These results indicated that the reconstituted 67 kD product comprised of components from both NCre-IntN and IntC-CCre. 1(E) Design of the *in vitro* reporter assay. HEK293T cells were transfected with NCre-IntN or IntC-CCre alone or together with a LoxP-Stop-LoxP-DsRed reporter. 1(F) When NCre-IntN or IntC-CCre were transfected alone, no DsRed expression was observed. When NCre-IntN and IntC-CCre were cotransfected along with the reporter in HEK293T cells, we observed strong DsRed expression. Scale bar, 50 μm.

Next, to validate whether the reconstituted Cre recombinase via trans-splicing of split inteins was functional, we attached fluorescent tags, mTagBFP2 and Cerulean to NCre-IntN and IntC-CCre respectively, to visualize the cellular expression of these constructs in HEK293T cells (Fig. 1E). We reasoned that the expression of these split-Cre constructs together with a LoxP-stop-LoxP DsRed reporter cDNA would easily indicate the activity and efficiency of split-Cre recombination through the presence of DsRed fluorescence in cells (Fig. 1E). As expected, we did not observe any DsRed fluorescence when either NCre-IntN or IntC-CCre was transfected alone. However, when both NCre-IntN and IntC-CCre were cotransfected, we observed strong DsRed signals, which was similar to cells transfected with the full-length Cre recombinase cDNA and reporter (Fig. 1F). We also observed that approximately 99% of the cells expressing both NCre-IntN and IntC-CCre exhibited DsRed fluorescence (Fig. 1F). Alternatively, we performed a luciferase assay as a readout for Cre-mediated recombination activity (Kaczmarczyk and Green, 2003) by transfecting either NCre-IntN or IntC-CCre alone or together into HEK293T cells expressing a LoxP-stop-LoxP luciferase cDNA construct (Fig. 2A). Twenty-four hours after transfection, we observed luminescence in only the cells cotransfected with both NCre-IntN and IntC-CCre (*p* < 0.001; Fig. 2B). Importantly, we observed that the Cre activities of reconstituted split intein split-Cre recombinase were comparable to those of native Cre recombinase (*p* = 0.798, no statistical significance; Fig. 2B (Ho et al., 2019)).

**Figure 2.**
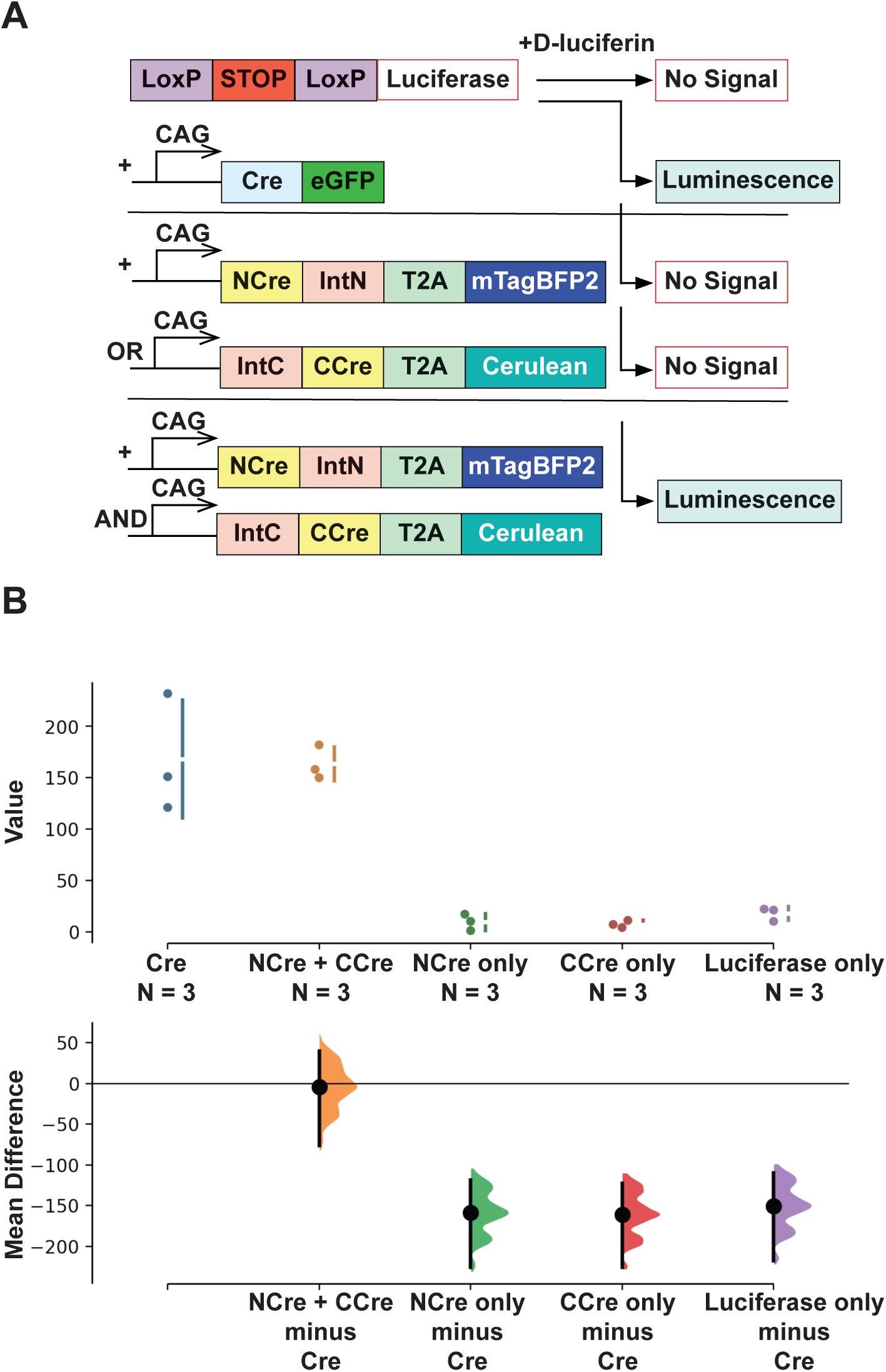
Design of luciferase assay. HEK293T cells were transfected with Cre recombinase, NCre-IntN or IntC-CCre alone, or together, with a LoxP-stop-LoxP-Luciferase reporter. 2(A) Schematic showing the experimental design for transfecting cDNA constructs in HEK293T cells. 2(B) The Cre activity is defined by the amount of luminescence detected in the luciferase assay. The mean difference for 4 comparisons against the shared control Cre recombinase are shown in the above Cumming estimation plot. The raw data is plotted on the upper axes. On the lower axes, mean differences are plotted as bootstrap sampling distributions. Each mean difference is depicted as a dot. Each 95% confidence interval is indicated by the ends of the vertical error bars. Importantly, we observed similar levels of Cre activity for both the native Cre recombinase and the reconstituted split-Cre recombinase. * *p* < 0.05, ** *p* < 0.01, *** *p* < 0.001.

To test if the split-Cre system is functional in the *in vivo* mammalian brain, we delivered NCre-IntN and IntC-CCre, Cre recombinase, or GFP only constructs into the lateral ventricles of LoxP-stop-LoxP-TdTomato reporter mice on embryonic day 16.5 (E16.5) via *in utero* electroporation (Zhang et al., 2016) (Fig. 3A). We did not observe any TdTomato expression when we electroporated GFP only or IntC-CCre fragment only (Fig. 3B). However, we observed strong TdTomato fluorescence in neurons when we electroporated both NCre-IntN and IntC-CCre constructs, and the fluorescence signal as well as a number of expressed neurons were similar to electroporation of native Cre recombinase (Fig. 3B). Taken together, these data showed that the reconstituted split intein-mediated split-Cre recombinase is functional in both *in vitro* and *in vivo* in mammalian brains.

**Figure 3.**
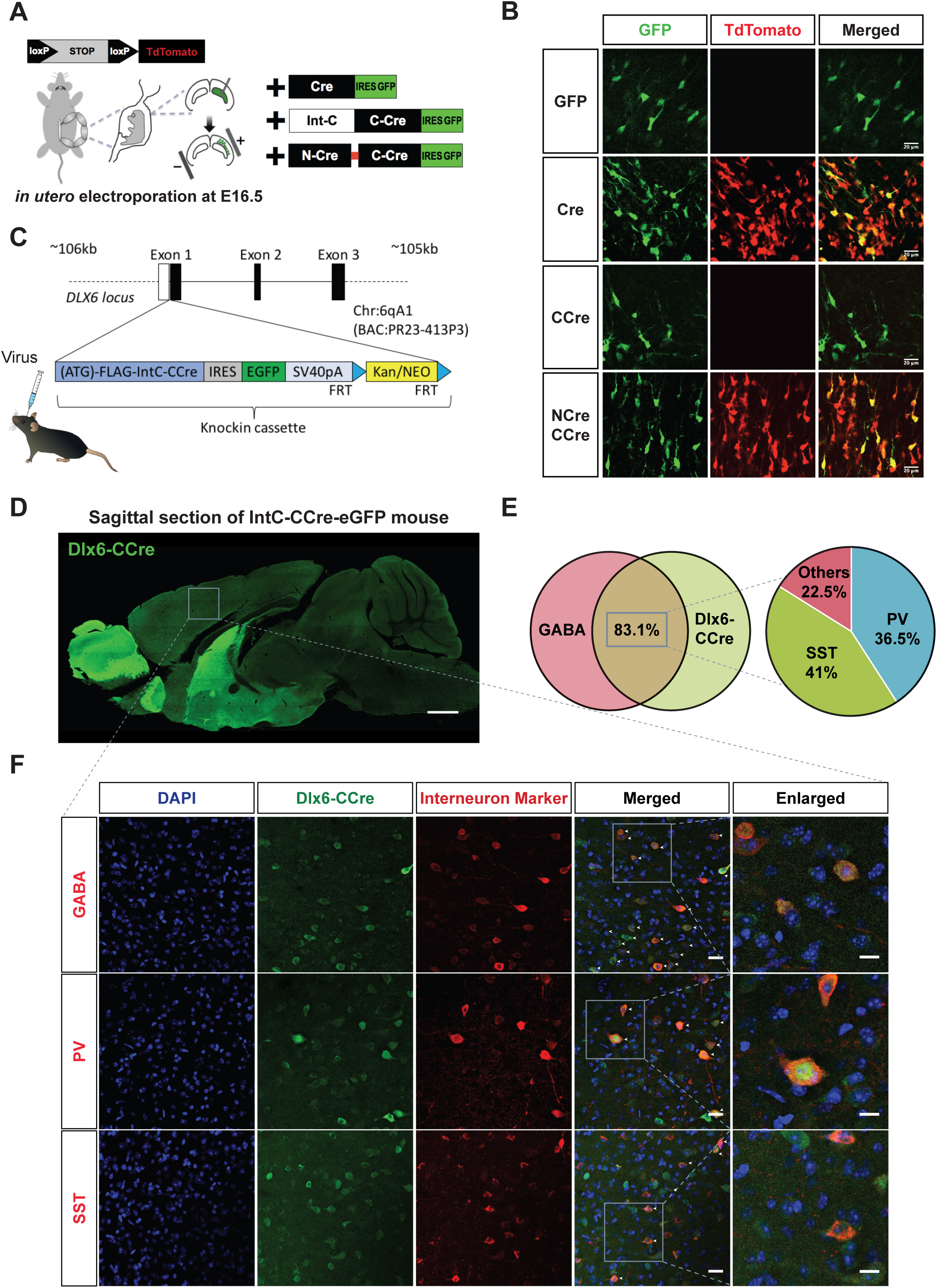
*In vivo* validation of the split intein-mediated split-Cre recombinase system. 3(A) Schematic showing the experimental design for electroporating constructs into LoxP-Stop-LoxP-TdTomato E16.5 embryos. 3(B) Strong TdTomato expression was observed with native Cre recombinase or when both NCre-IntN and IntC-CCre constructs were electroporated. No TdTomato was observed when only IntC-CCre was electroporated alone. This demonstrated that Cre-mediated recombination occurs only when both NCre and CCre are present. 3(C) Schematic diagram of BAC transgenic mice expressing IntC-CCre in forebrain GABAergic neurons. The resultant BAC transgenic mouse was then further crossed with the LoxP-Stop-LoxP-TdTomato reporter line mouse. Scale bar, 20 μm. 3(D) Sagittal section from a DLX-CCre-IRES-eGFP mouse showing that IntC-CCre-positive cells were predominantly found in the forebrain. Scale bar, 1 mm. 3(E) Venn diagram showing that 83.1% of DLX6-CCre neurons were GABA-positive. Of the CCre+/GABA+ neurons, 36.5% were PV and 41% were SST. 3(F) Immunohistochemical analyses showing DLX6-CCre-IRES-eGFP brain sections stained using specific antibodies against GABA, PV and SST. Scale bars, 20 μm (merged) and 10 μm (enlarged).

To target long-range GABAergic projection neurons (Tamamaki and Tomioka, 2010), we generated a transgenic mouse that expressed IntC-CCre-IRES-eGFP under the control of an endogenous *Dlx6* promoter known to be expressed exclusively in forebrain GABAergic neurons (de Lombares et al., 2019; Dimidschstein et al., 2016; Le et al., 2017; Wang, 2011) (Fig. 3C and D). Next, the resulting *Dlx6*-CCre-IRES-eGFP mice were further crossed with Rosa26-LoxP-stop-LoxP-TdTomato reporter mice (Madisen et al., 2010) (Fig. 3D). Using immunohistochemistry, we confirmed that IntC-CCre expression was restricted to GABAergic neuronal populations (Fig. 3E and F). A total of 83.1% of the IntC-CCre neurons, indicated by green fluorescence, expressed GABA, and 40.9% and 43% of these neurons expressed PV and SST respectively (Fig. 3E and F).

First, we validated the reconstitution of split-Cre in our *Dlx6*-CCre-IRES-eGFP mice. We generated lentiviral vectors encoding either mTagBFP2 or NCre-IntN-mTagBFP2. We injected mTagBFP2 lentiviral particles as a control into one hemisphere of the striatum and injected NCre-IntN-mTagBFP2 particles into the other (Fig. 4A). While we did not observe any TdTomato expression in the hemisphere injected with mTagBFP2, robust TdTomato expression was evident in the hemisphere injected with NCre-IntN-mTagBFP2 (Fig. 4B). These results indicated that selective reconstitution of recombinatory Cre function occurred only when both NCre-IntN and IntC-CCre were expressed. Of the 37 neurons expressing both NCre-IntN and IntC-CCre, 78.3% exhibited a strong TdTomato fluorescence signal (Fig. 4C).

**Figure 4.**
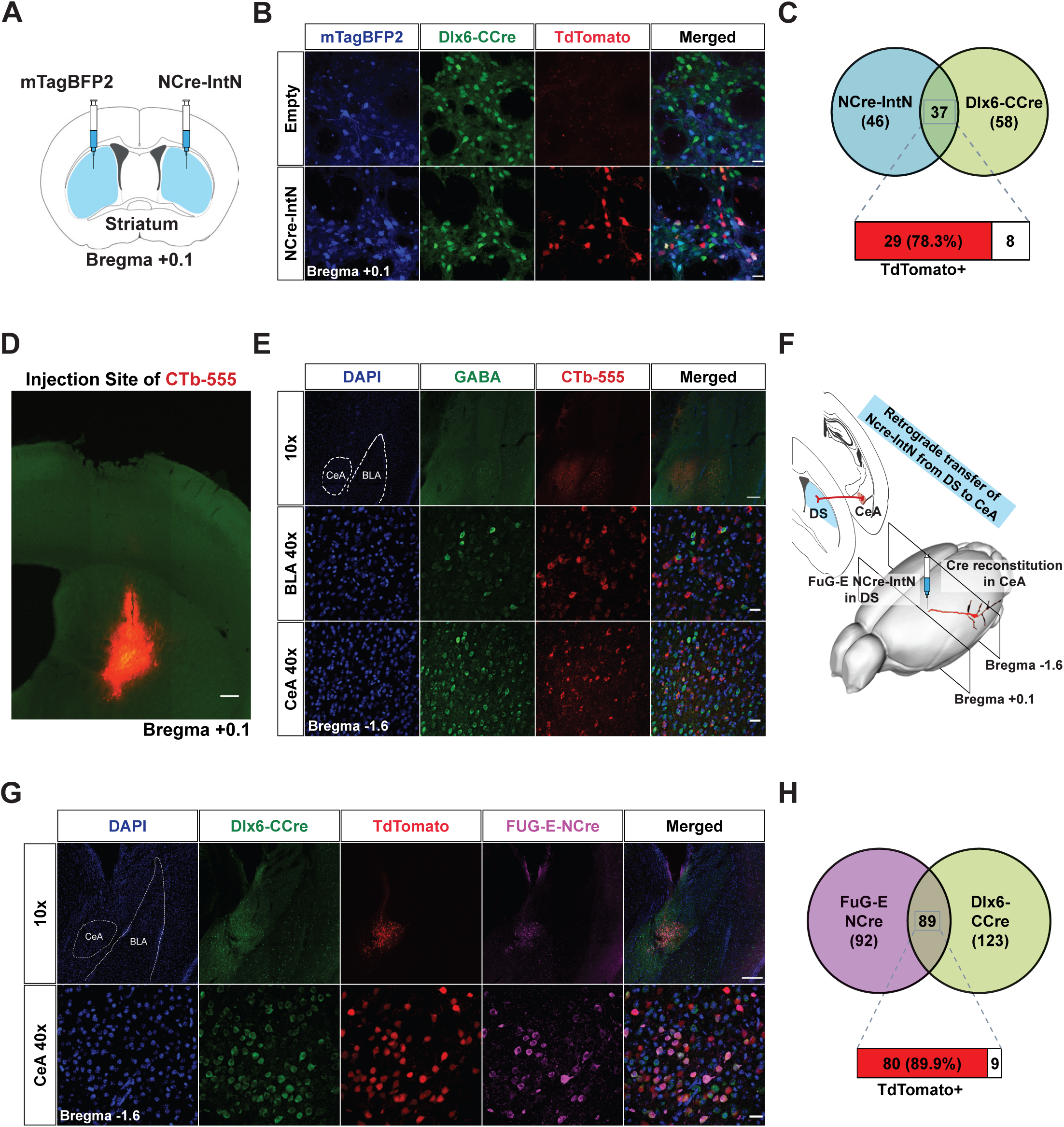
Targeting long-range GABAergic neurons using a DLX6-CCre-IRES-eGFP mouse and virus. 4(A) Schematic of stereotaxic injection of lentiviral particles to observe recombination of Cre recombinase in DLX6-CCre-IRES-eGFP mice. The mTagBFP2-only lentiviral particles were injected into one hemisphere or the striatum, while the NCre-IntN-carrying lentiviral particles were injected into the other hemisphere. 4(B) In the hemisphere injected with the mTagBFP2-only lentiviral particles, no TdTomato signal was observed. However, in the hemisphere injected with NCre-IntN-carrying lentiviral particles, a TdTomato signal was observed, indicating the presence of functional Cre recombinase. Scale bar, 25 μm. 4(C) Venn diagram showing the percentage of cells expressing TdTomato. 4(D) Microscopic image showing the injection site of CTb-555 in the DS. Scale bar, 200 μm. 4(E) CTb-555-labeled cells were observed in both the CeA and BLA. However, immunohistochemical analysis illustrated that only labeled cells in the CeA were GABAergic, while those in the BLA were not. Scale bars, 200 μm (10x) and 20 μm (40x). 4(F) Schematic diagram showing projection from CeA to DS. The FuG-E-NCre virus was injected into the DS, which was retrogradely transferred to the CeA. Cre recombination is expected to occur in GABAergic neurons in the CeA that project to the DS. 4(G) When FuG-E-NCre pseudotyped lentiviral particles were injected into the DS of DLX6-CCre-IRES-eGFP mice crossed with the reporter line, TdTomato cells were observed in the CeA. Scale bars, 200 μm (10x) and 20 μm (40x). 4(H) Venn diagram showing the percentage of cells expressing TdTomato in the CeA.

To selectively target long-range GABAergic projection neurons, we first validated the presence of these types of neurons that were found to project from the central amygdala (CeA) to the dorsal striatum (DS) (Lingawi and Balleine, 2012) by injecting cholera toxin subunit B fused to Alexa Fluor 555 (CTb-555) into the DS of wild-type mice (Fig. 4D) (Vercelli et al., 2000). We observed CTb-555 labeling in both the CeA and basolateral amygdala (BLA) (Fig. 4E). Immunohistochemistry analyses showed that only CTb-labeled neurons in the CeA were GABA-positive, whereas the CTb-labeled neurons in the BLA were not (Fig. 4E). This confirmed that only the projections from the CeA to the DS were from GABAergic neurons, as reported by Lingawi and Balleine (2012). Next, we generated retrograde lentiviral particles carrying NCre-IntN by substituting the vesicular stomatitis virus glycoprotein (VSV-G) with FuG-E, a pseudotyped lentiviral envelope where its fusion glycoprotein comprises of VSV-G and the rabies virus glycoprotein (RV-G) (Kato et al., 2014). This allows NCre-IntN to be retrogradely transferred from the axon terminating in the DS back to the cell body of neurons in the CeA (Fig. 4F). Injection of the FuG-E-NCre-IntN viral particles into the DS of the *Dlx6*-CCre-IRES-eGFP::LoxP-stop-LoxP-TdTomato mice resulted in TdTomato-expressing neurons within the CeA (89.9% NCre- and CCre-expressing neurons expressed TdTomato)(Fig. 4G and H). While we observed NCre-IntN expression in some BLA neurons as the BLA also projects to the DS, we did not observe a TdTomato signal, indicating that Cre reconstitution was highly specific (Fig. 4G).

## Discussion

Here, we developed a split-Cre system based on split intein trans-splicing mechanisms. By generating a transgenic mouse model that expresses IntC-CCre in forebrain GABAergic neurons and NCre-IntN expressing retrograde viral particles, we could selectively target long-range GABAergic neurons projecting from the CeA to the DS (Fig. 4G). We are still puzzled by the slightly lower recombination efficiency of our split-Cre system upon viral delivery schemes *in vivo* (approximately 80%, compared to 99% upon transfection; Fig. 4C and H), which could simply be due to the low titer (3E8 IU/mL) of our lentiviral particles. However, this result also indicates the selectivity and stringency of our split-Cre recombinase, since germline recombination and transient unwanted expression of Cre recombinase have been reported in even well-established transgenic mouse lines (Song and Palmiter, 2018). Given that we can genetically target long-range GABAergic neurons with high specificity, we can further investigate the functions of long-range GABAergic projection neurons during active behavior using existing Cre-dependent opto- and chemogenetic tools. In addition, our approach can be used to understand whether the projection patterns of a neuron correlates with its gene expression profile using RNA sequencing. Our approach should be able overcome the limitation of having few virally packageable, tissue-specific promoters (Nathanson et al., 2009).

Taken together, our combinatory split intein-mediated split-Cre system allows us to limit Cre recombinase expression to specific subsets of neurons by means of its gene expression profile and anatomical connections, enabling the systematic study of specific types of neurons in the mammalian brain.

## Methods

### Computer simulation of split-intein split-Cre recombinase product

Model for split-intein mediated split-Cre recombinase-LoxP DNA complex was calculated using automated modelling mode in the SWISS model server (http://swissmodel.expasy.org/), based on the crystal structure of Cre-LoxP synaptic complex (1NZB) as a template (Ennifar et al., 2003). Split inteins is expected to form stable β-turn structure without any substantial structural changes in the original Cre recombinase. Figures for the structure were prepared using PyMol software (DeLano, 2002).

### Construction of NCre-IntN and IntC-CCre

NCre and CCre were generated from the pCAG-Cre:GFP, a gift from Connie Cepko (Addgene plasmid #13776; https://www.addgene.org/13776/; RRID:Addgene_13776) using the KAPA HiFi PCR kit (Roche Sequencing). IntN and IntC were amplified from the Npu DnaE inteins with site directed mutagenesis according to Lockless and Muir (2009). All plasmids were subcloned into a pCDH expression vector (System Biosciences), assembled using Gibson Assembly (New England Biolabs) and verified by Sanger DNA sequencing.

### Western blot analyses

HEK293T cells in 12-well culture plates were transfected with 1ug of each construct and Lipofectamine 2000 (Thermo Fisher Scientific) for 18 hours. Cells were supplied with fresh media after transfection and grown for another 2 days. Cells were mechanically lysed in RIPA buffer containing phosphatase and protease inhibitors. The proteins were separated by SDS-PAGE under reducing conditions and transferred to polyvinyl difluoride (PVDF) membranes (Millipore, Billerica, MA). Antibody incubations were performed in 3% BSA in TBS buffer using the following antibodies: anti-HA, 1:1000 (MilliporeSigma); anti-FLAG, 1:1000 (MilliporeSigma); anti-β-Tubulin, 1:1000 (Covance).

### HEK293T in vitro reporter assay

HEK293T cells were seeded into 12-well culture plates containing coverslips treated with poly-L-lysine and transfected the next day with 1μg of each construct using Lipofectamine 2000 (ThermoFisher Scientific) for 18 hours. The cells were supplied with fresh media after transfection and subsequently fixed with 4% (w/v) paraformaldehyde in PB 24 hours later. The coverslips were mounted onto microscope slides with FluorSave mounting medium (Calbiochem) and imaged with the Zeiss LSM780 confocal microscope (Zeiss Microscopy, Munich, Germany) using the 40x oil objective lens.

### HEK293T luciferase assay

HEK293T cells were seeded into 96-well culture plates and transfected with 1g of each constructed the next day. 24 hours post transfected, we washed the cells with PBS and treated them with lysis buffer (Pierce Firefly Luciferase Glow Assay Kit, Thermo Fisher Scientific) for 20 minutes. The cell lysate was then collected and spun down to remove cell debris. 15µl of the cell lysate was transferred into a 96-well black polysterene plate (Corning Costar). 45µl of the working buffer (D-luciferin and the Firefly Glow Assay buffer according to manufacturer’s instructions, Pierce Firefly Luciferase Glow Assay Kit, Thermo Fisher Scientific) was added to the cell lysate and we waited 10 minutes before acquiring signals using a microplate reader (Tecan, Männedorf, Switzerland).

### Animals

Animals were housed in a specific pathogen-free facility maintained below 22°C and 55% humidity under a 12h light-dark cycle (lights on at 0700h) with *ad libitum* access to food and water. All experimental procedures were conducted in accordance with guidelines for the care and use of laboratory animals for scientific purposes with approved protocols from the Institution Animal Care and Use Committee of SingHealth Research (Protocol number: 2014/SHS/921).

### In utero electroporation

Floxed Tdtomato transgenic females were time mated. On embryonic day 16.5 (E16.5), *ex vivo* electroporation and organotypic brain slice culture was performed as described elsewhere (Lizarraga et al., 2012). A GFP expression plasmid was used as a negative control, and a Cre plasmid was used as a positive control. For the experiments embryos were electroporated either with NCre plasmid only or with both NCre and CCre plasmids. For image acquisition, on day 3 of culture, the organotypic slice inserts were transferred to a 50mm petri dish containing culture medium and imaged using an inverted long distance fluorescent microscope.

### Generation of DLX6-IntC-CCre-eGFP mice

We used the bacterial artificial chromosome (BAC) transgenic approach to generate the animals that express IntC-CCre under the DLX6 promoter. In this experiment, we constructed a knock-in cassette containing a FLAG-tagged (at the N-terminal of) IntC-CCre coding sequence followed by an IRES-eGFP cassette. A SV40 polyadenylation signal sequence was placed at the 3’ end of the cassette to stabilize the transcript. A FRT flanked Kan/Neo selection cassette was attached to the 3’ end of the IntC-CCre cassette to facilitate screening of the recombinant clones. We used recombineering technology to integrate the IntC-CCre expression cassette into the BAC clones containing the Dlx gene locus. In the recombinant BAC clones, the translation start codon ATG of FLAG-IntC-CCre is fused to the ATG of the endogenous Dlx gene. After deletion of the selection cassette by expression of Flp, the resultant BAC clones containing the IntC-CCre cassette were used to create the transgenic mice by standard pronuclear injection.

### Virus preparation

HEK293T cells were cultured in T75 flasks and transfected with hSyn.NCre-IntN.T2A.mTagBFP2, envelope and VSV-G or FUG-E packaging plasmid. pCAGGS-FuG-E was a gift from Dr. Kazuto Kobayashi (Addgene plasmid #67509; http://n2t.net/addgene:67509; RRID:Addgene_67509). 18 hours after transfection, we supplied fresh medium and the cells were further incubated for 48 hours. The medium was collected and filtered through a 0.45μm Minisart syringe filter (Sartorius Stedim, Goettingen, Germany). Viral vector particles were pelleted via ultracentrifugation at 25,000g for 2 hours. Supernatant was subsequently removed and the pellet was resuspended in sterile PBS overnight. Virus titre was quantified using the qPCR Lentivirus Titration Kit according to manufacturer’s directions (Applied Biological Materials, British Columbia, Canada).

### Stereotaxic injection of virus and cholera toxin subunit B

Mice were anesthetized with 2% isoflurane in oxygen at a flow rate of 0.5L/min. Mice were placed in a stereotaxic apparatus to receive bilateral infusions of the virus or 0.5% (w/v) cholera toxin subunit B (CTb) Alexa Fluor 555 Conjugate (Invitrogen) into the dorsal striatum (+0.1mm antero-posterior, 2.0mm mediolateral, 2.7mm dorsoventral); coordinates relative to bregma (Paxinos and Watson, 2007). 1ul of virus or 0.1μl of CTb-555 was infused per hemisphere with a 30-gauge needle (Hamilton, United States) over 5 minutes and 0.5 minute respectively using a programmable infusion pump (Hamilton, United States). The needle was left in the place for 5 minutes to allow diffusion of the virus or CTb before removal at 0.1mm/sec. Following surgery, mice received 5mg/kg carprofen intraperitoneally. Mice were sacrificed 28 days or 7 days following virus or CTb injection respectively.

### Perfusion and immunohistochemical analyses

Mice were anesthetized with isoflurane and transcardially perfused with ice-cold PBS, followed by 4% (w/v) paraformaldehyde (PFA) in 0.1M PB. Extracted brains were submerged in PFA for an hour before transferring to 30% (w/v) sucrose in 0.1M PB for cryoprotection. After 48 hours, brain were sectioned at 40um and stored in PBS-Azide. Sections were first permeabilized in 0.2% Triton-X100 in PBS (30min, RT) and later incubated in (i) a blocking solution of in PBS containing 5% donkey serum, 3% bovine serum albumin and 0.2% Triton-X100 (1h, RT); (ii) the same solution containing the primary antibody overnight at 4°C (anti-tRFP, 1:1000, Evrogen; anti-GABA, 1:2000, MilliporeSigma anti-parvalbumin, 1:2000, Abcam; anti-somatostatin, 1:1000, Peninsula Laboratories) (iii) the same solution containing a 1:500 dilution of secondary antibody (2h, RT). All sections were then stained with DAPI (1:5000 diluted in 0.2% Triton X-100 in PBS, Sigma-Aldrich) and mounted with FluorSave (Merck Millipore). Images of the immunostained sections were acquired using the widefield (Nikon Instruments, Tokyo, Japan) or confocal microscope (Zeiss Microscopy, Munich, Germany).

### Statistical analyses

Data were analysed and plotted using Estimation stats/plots (Ho et al., 2019). *p* < 0.05 was considered statistically significant.

## List of abbreviations

BAC: bacterial artificial chromosomes
BLA: basolateral amygdala
CCre: C-terminal Cre fragment
CeA: central amygdala
CTb: cholera toxin subunit B
DNA: deoxyribonucleic acid
DS: dorsal striatum
FuG-E: fusion glycoprotein type E
GABA: gamma aminobutyric acid
HEK293T: human embryonic kidney 293 cells expressing mutant version of SV40 large T antigen
IntC: C-terminal intein
IntN: N-terminal intein
NCre: N-terminal Cre fragment
PV: parvalbumin
RNA: ribonucleic acid
RV-G: rabies virus glycoprotein
SST: somatostatin
VSV-G: vesicular stomatitis virus glycoprotein

## Acknowledgments

We would like to thank Dr. Henry Yin, Dr Qing Yan, and George Yu for experimental assistance. We would like to also thank Dr. Wang Linfa and Dr. Danielle Anderson for their advice and help with viral constructs.

## Author contributions

A.K. and H.S.J. constructed plasmids. H.K. generated the 3D model of reconstituted split-intein mediated split-Cre recombinase. H.S.J. performed western blot analyses. A.K. designed and performed HEK293T reporter assays, *in vivo* stereotaxic injections, genotyping and histological analyses. P.K. performed *in utero* electroporation. H.S.J. supervised project. A.K. and H.S.J. contributed to ideas, designed and wrote manuscript.

